# Specific *gyrA* gene mutations correlate with high prevalence of discordant levofloxacin resistance in *Mycobacterium tuberculosis* isolates from Beijing, China

**DOI:** 10.1101/841031

**Authors:** Fengmin Huo, Yifeng Ma, Shanshan Li, Yi Xue, Yuanyuan Shang, Lingling Dong, Yunxu Li, Yu Pang

**Affiliations:** National Clinical Laboratory on Tuberculosis, Beijing Key laboratory on Drug-resistant Tuberculosis Research, Beijing Chest Hospital, Capital Medical University, Beijing Tuberculosis and Thoracic Tumor Institute, Beijing, China

**Keywords:** tuberculosis, levofloxacin, drug susceptibility testing, MeltPro assay, *gyrA*

## Abstract

Although molecular diagnostics are highly sensitive in the diagnosis of fluroquinolone (FQ)-resistant tuberculosis (TB), the discordant results with the sequential phenotypic drug susceptibility testing (DST) invariably occur. To investigate the prevalence and the cause of discordant results between molecular and phenotypic DST for levofloxacin (LFX), the cases who were determined to be LFX-susceptible using the phenotypic DST but LFX-resistant using the MeltPro assay were retrospectively reviewed in Beijing, China. We measured LFX minimal inhibitory concentrations (MICs) using Middlebrook 7H9 broth. Sanger sequencing was used to determine genotypic characteristics of discordant isolates. Between January and December 2018, 126 (22.1%) out of 571 smear-positive TB patients were identified as LFX-resistant TB by MeltPro assay. Among the 126 LFX-resistant TB, there were 34 isolates identified as LFX-susceptible TB by phenotypical DST. This result demonstrated a discordance prevalence of 27.0%. LFX MICs were majorly centered around the critical concentration, and 7 (21.2%) and 13 (39.4%) had MICs of 2.0 mg/l and 4.0 mg/l, respectively. The most prevalent mutations conferring discordant LFX resistance were the amino acid substitutions of Ser to Leu in 90 codon (13, 39.4%) and Asp to Ala in 94 codon of *gyrA* (11, 33.3mg/l belonged to Ala90Val and Asp94Ala group. In conclusion, our data demonstrate that more than one quarter of LFX-resistant isolates by molecular method are susceptible to LFX using the phenotypic DST. The high prevalence of discordant LFX resistance is majorly due to the isolates with specific mutations within *gyrA* gene conferring proximity of MICs to the critical concentration.

## INTRODUCTION

Despite the great achievements in tuberculosis (TB) control globally, the epidemic of drug-resistant tuberculosis, especially multidrug-resistant tuberculosis (MDR-TB), poses a major barrier to eliminate TB (1, 2). In 2018, the World Health Organization (WHO) estimates that 48, 400 MDR/RR-TB cases have occurred worldwide, only 32% of which were successfully diagnosed and treated (2). Therefore, rapid and accurate laboratory diagnosis of drug susceptibility of MTB is essential to ensure early initiation of appropriate treatment for drug-resistant TB patients, thus preventing further transmission in the community (3, 4).

Conventional diagnosis of drug-resistant TB requires the phenotypical drug susceptibility testing (DST) using growth-based methods (5). Due to the slow growth rate of tubercle bacilli, it takes weeks to months to yield final results (5). Recently, several commercial molecular tests have been endorsed by WHO for the initial diagnosis of drug-resistant TB, such as GeneXpert MTB/RIF (Cephid, USA), and GenoType MTBDRplus (Hain Lifescience, Germany) (6, 7). These tests detect drug resistance by identifying the core genetic regions responsible for drug resistance in a timely manner (3). Although a serial of clinical trials have demonstrated that molecular tests are highly sensitive in the diagnosis of drug-resistant TB, especially for rifampin, isoniazid and fluroquinolone, the discordant results with the sequential phenotypic DST invariably occur (8). The situation is more complicated if the “false-positive” result is reported by molecular test considering that the patients have undergone the therapy for drug-resistant TB. Discordant results between molecular and phenotypic rifampicin susceptibility testing are reported, resulting in diagnostic and clinical management dilemmas (9). Understanding the interpretation of these discordant results is thus important to ensure optimal diagnosis and treatment.

Fluroquinolones (FQs) are cornerstone drugs in the treatment of MDR-TB (10). The subunit A of gyrase (*gyrA*) is the primary target of current FQs in MTB, thus killing bacteria via inhibiting DNA supercoiling and disrupting DNA replication (11, 12). Thus, the mutations located in *gyrA* gene are considered as an alternative for predicting FQ resistance in MTB (12). In China, the MeltPro TB assay (MeltPro, Zenssen, China) is the only commercial molecular test approved for the diagnosis of FQ-resistant TB, and also widely used in clinical practice since 2018 (13). However, limited knowledge is available to examined the prevalence of false-positive results between molecular and phenotypic DST for FQs. To address this concern, we retrospectively reviewed the cases who were determined to be levofloxacin (LFX)-susceptible using the phenotypic DST (pDST) but LFX-resistant using the MeltPro assay.

## METERIALS AND METHODS

### Study Design

This is a retrospective analysis of laboratory results and stored clinical isolates were collected for routine clinical practice. The study was conducted in the Beijing Chest Hospital, which is the National Clinical Centre on Tuberculosis in China. Consecutive individuals with LFX resistance by MeltPro were reviewed between January 2018 and December 2018. The cases without confirmatory LFX resistance testing were excluded from further analysis. We collected the MTB isolates from the cases with discordant LFX resistance results between MeltPro and phenotypic DST. Discordance was defined as an initial MeltPro result with LFX resistance with only LFX-susceptible follow-up phenotypic DST tests. The sputum specimens from the smear-positive TB patients were collected for MeltPro TB assay. The MeltPro assay testing was performed as previously described (Fig. 1) (13). In addition, the commercial 96-well plate containing lyophilized antibiotics were used for determination of *in vitro* susceptibility of MTB isolates to first- and second-line anti-tuberculosis drugs according to the manufacturer’s instructions (Encode Medical Engineering Co., Zhuhai, China). Susceptibility was judged on the basis of the following drugs and concentrations: isoniazid (0.2 mg/l), rifampicin(1.0 mg/l), levofloxacin(2.0 mg/l), ethambutol(2.5 mg/l), amikacin(1.0 mg/l), capreomycin(2.5 mg/l).

**Figure 1.**
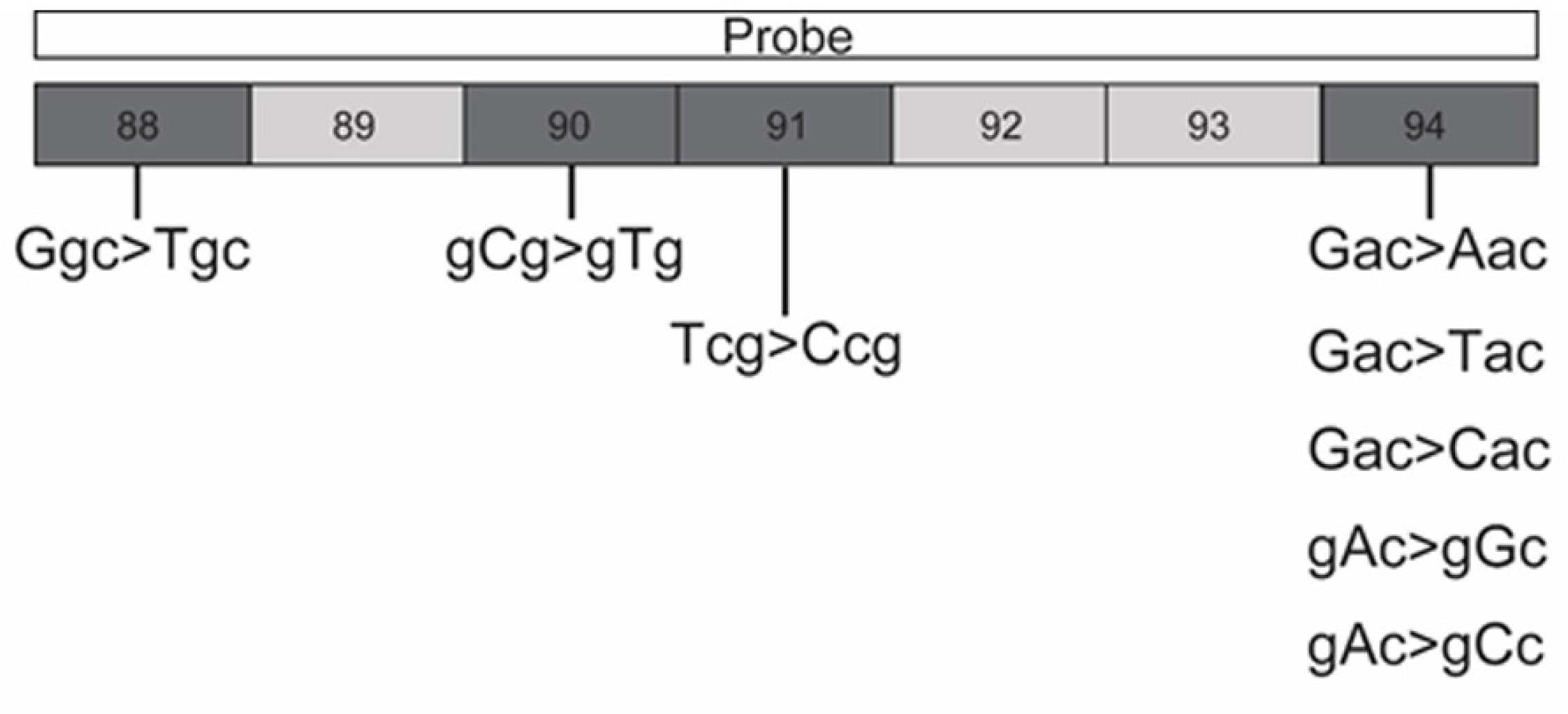
The region of *gyrA* targeted by the probe of MeltPro TB assay. A specific probe covers region of *gyrA* codons 88 to 94, consisting of the most common mutation types conferring FQ resistance. The probe is perfectly matched with the wild-type sequence of *gyrA* gene. The corresponding genotype was interpreted according to the T_m_ of melting curve of double-stranded structures between probe and targeted *gyrA* gene.

### Bacterial strains and culture condition

All MTB isolates were stored in the Middlebrook 7H9 medium containing 5% glycerol at the BioBank of Tuberculosis, Beijing Chest Hospital. Prior to experiments, the bacterial cells were recovered on Lowenstein-Jensen (L-J) medium for 4 weeks at 37°C.

### Minimal inhibitory concentration (MIC)

The MICs for LFX were determined in Middlebrook 7H9 broth containing oleic acid-albumin-dextrose-catalase (OADC) (14). Fresh colonies were harvested from the surface of L-J medium, and added to a saline solution with glass beads. After vortex for 1 min, the suspension was adjusted to turbidity at a McFarland standard of 1.0. 200 μl bacterial diluent was transferred to 20 ml of 7H9 broth, and 100 μl of this dilution was inoculated into each well. The drug concentrations tested were doubling dilution, ranging from 0.063 to 64 mg/l. Plates were incubated at 37°C in 5% CO_2_ for one week. Then 70 μl of AlamarBlue and 5% Tween80 2:5 mixed solution was added to each well, incubated for 24 h at 37°C, and assessed for color development. As a consequence of bacterial growth, the oxidation-reduction indicator AlamarBlue changed color from blue to pink (15). The MIC was defined as the lowest concentration of drug that prevented the color change from blue to pink. The reference MTB strain H37Rv (ATCC 27249) was tested in both rounds for quality control purpose.

### DNA sequencing

The crude genomic DNA was extracted from freshly cultured bacteria using the simple boiling method as previously described (16). The genomic DNA was used as template to conduct PCR amplification. The partial fragment of *gyrA* gene, containing quinolone resistance-determining region, were amplified (17). The PCR mixture were prepared as follows: 25 μl 2×PCR Mixture, 2 μl of DNA template and 0.2 μM of each primer set. After completion of the thermal cycler program, the amplicons were sent to Tsingke Company (Beijing, China) for DNA sequencing service. DNA sequences were compared with the homologous sequences of MTB H37Rv strain to identify whether the sequences had genetic mutations within QRDR.

## RESULTS

Between January and December 2018, 571 smear-positive TB patients were screened by MeltPro assay (Fig. 2). Of these, 126 (22.1%) were identified as LFX-resistant TB. Phenotypical DST showed that 92 isolates were LFX-resistant, while the remaining 34 were LFX-susceptible, demonstrating a discordance prevalence of 27.0%. After exclusion of one isolate due to subculture failure, a total of 33 discordant isolates from 33 patients were further analyzed. As showed in Table 1, the isolates displayed a wide range of MICs with the majority centered around the critical concentration of 2.0 mg/l. Out of isolates tested, 7 (21.2%) and 13 (39.4%) had MICs of 2.0 mg/l and 4.0 mg/l, respectively.

**Figure 2.**
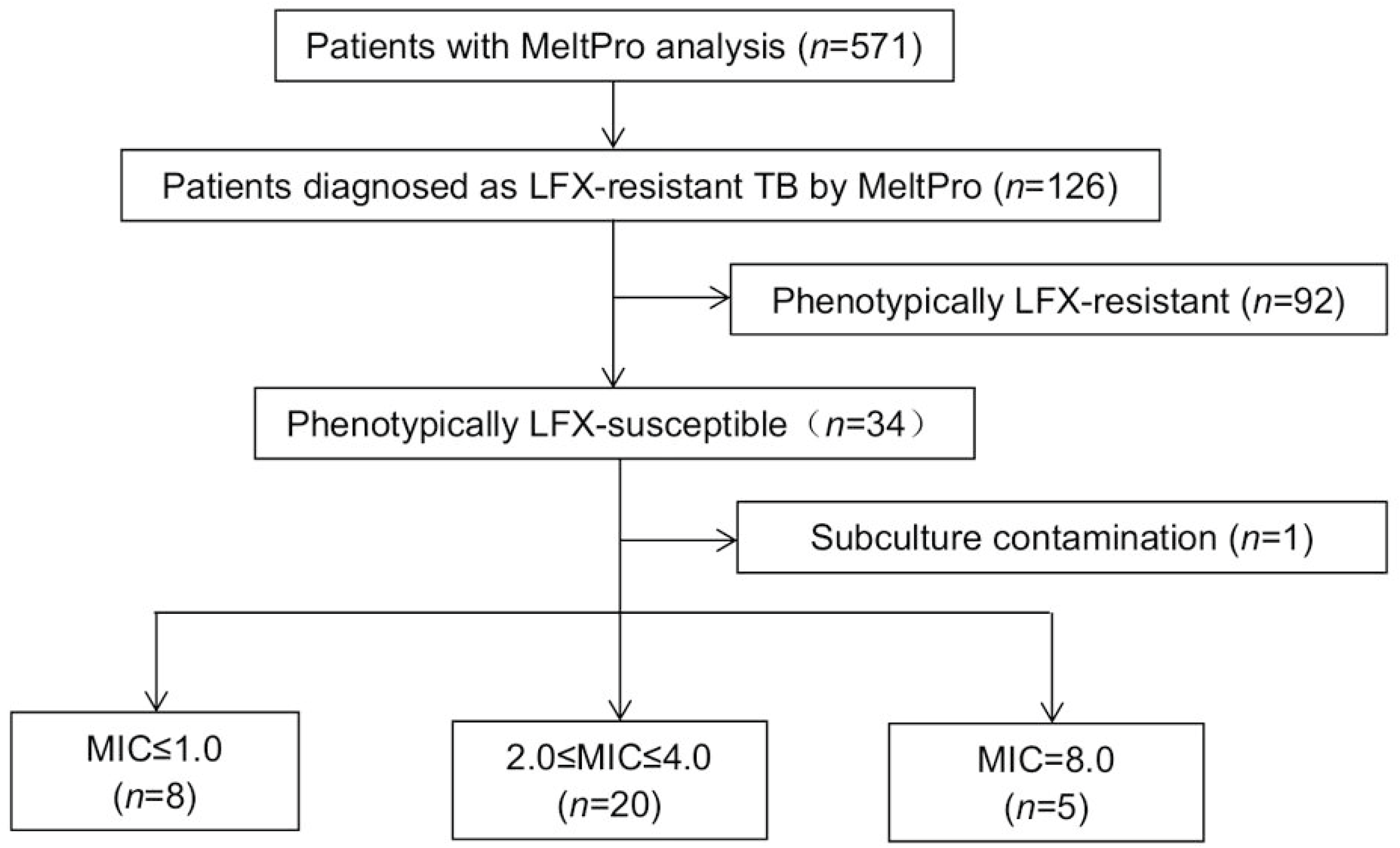
Enrolment of MTB isolates in this study.

**Table 1.** Distribution of MTB isolates with different MICs stratified by *gyrA* mutations

| Mutation type | No. of isolates with different MICs (mg/L) |  |  |  |  | Total<br>(n=33) |
| --- | --- | --- | --- | --- | --- | --- |
|  | ≤0.5<br>(n=2) | 1<br>(n=6) | 2<br>(n=7) | 4<br>(n=13) | 8<br>(n=5) |  |
| Ala90Val | 0(0.0) | 0(0.0) | 4(57.1) | 7(53.8) | 2(40.0) | 13(39.4) |
| Thr91Pro | 0(0.0) | 0(0.0) | 0(0.0) | 0(0.0) | 1(20.0) | 1(3.0) |
| Asp94Ala | 0(0.0) | 6(100.0) | 3(42.9) | 2(15.4) | 0(0.0) | 11(33.4) |
| Asp94Gly | 0(0.0) | 0(0.0) | 0(0.0) | 0(0.0) | 1(20.0) | 1(3.0) |
| Asp94Val | 0(0.0) | 0(0.0) | 0(0.0) | 1(7.7) | 0(0.0) | 1(3.0) |
| Asp94Tyr | 0(0.0) | 0(0.0) | 0(0.0) | 1(7.7) | 0(0.0) | 1(3.0) |
| Asp94Asn | 0(0.0) | 0(0.0) | 0(0.0) | 0(0.0) | 1(20.0) | 1(3.0) |
| Heteroresistance<sup>a</sup> | 0(0.0) | 0(0.0) | 0(0.0) | 2(15.4) | 0(0.0) | 2(6.1) |
| Wide type | 2(100.0) | 0(0.0) | 0(0.0) | 0(0.0) | 0(0.0) | 2(6.1) |

<sup>a</sup>Heteroresistance was defined as a heterogeneous population of tubercle bacilli according to the sequencing chromatograms.

DNA sequencing data revealed that the most prevalent mutations conferring discordant FQ resistance were the amino acid substitutions of Ser to Leu in 90 codon (13, 39.4%) and Asp to Ala in 94 codon of *gyrA* (11, 33.3%), respectively. Mutations Thr91Pro *(n* =1), Asp94Gly (*n* =1), Asp94Val (*n* = 1), Asp94Tyr (*n* = 1) and Asp94Asn (*n* = 1) were detected in five separate isolates. In addition, two heteroresistant isolates were identified by Sanger sequencing plots. In one isolate, the mutant G allele in 94 codon of *gyrA* (94Gly) was mixed with the wild-type A allele (94Asp) (Fig. 3A). Double point mutations were observed in the other heteroresistant isolate carrying both the Thr91Pro and Asp94Gly mutations (Fig. 3B). Notably, two FQ-resistant isolates reported by MeltPro assay had no mutation within *gyrA* gene by DNA sequencing. By re-examination of MeltPro patterns, we found that both these isolates had two melting curve peaks (Fig. 4), representing the simultaneous presence of mutant and the wild type in the specimens.

**Figure 3.**
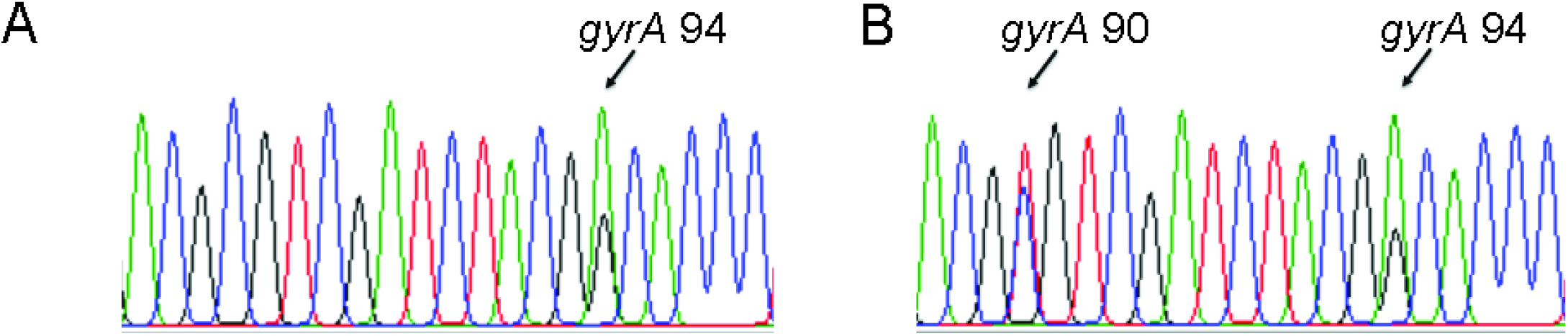
The *gyrA* gene sequencing chromatograms of two heteroresistant MTB isolates. A. The mutant G allele in 94 codon of *gyrA* (94Gly) was mixed with the wild-type A allele (94Asp). B. Double point mutations were observed in the heteroresistant isolate carrying both the Thr91Pro and Asp94Gly mutations.

**Figure 4.**
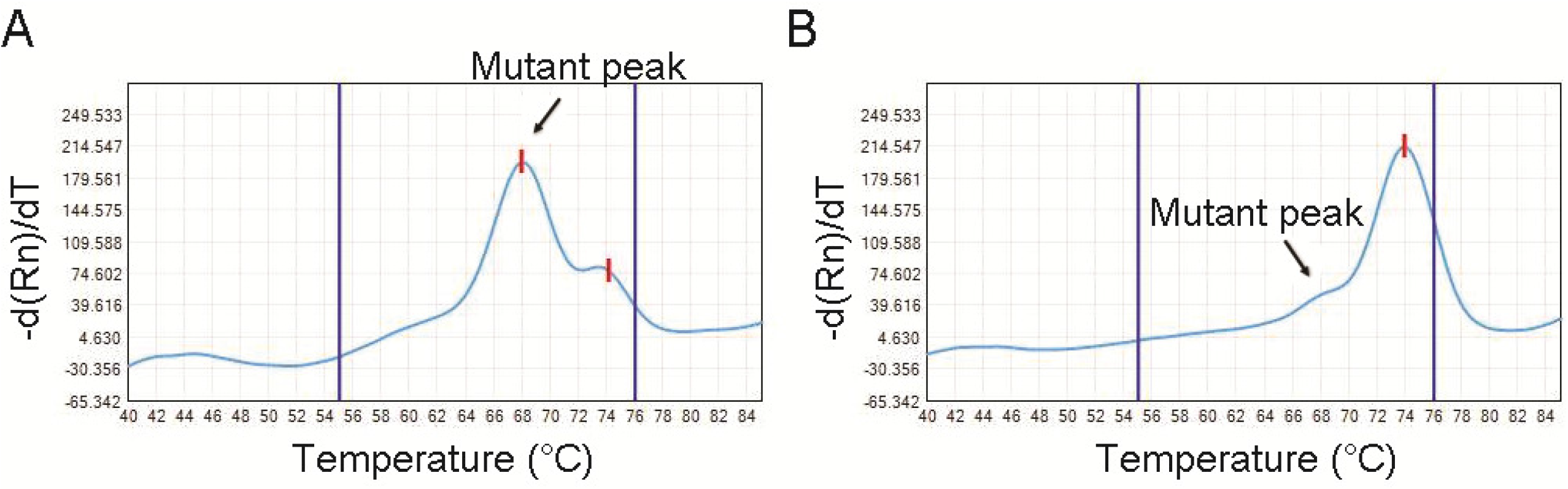
The raw melting curve analysis by MeltPro TB assay of two MTB isolates with wide-type *gyrA* locus. The two isolates both had two melting curve peaks in the melting curve, indicating the occurrence of wide-type and mutant *gyrA* gene in the initial specimens.

Mutations in the *gyrA* gene and corresponding MICs of the 33 selected isolates are shown in Table 2. 18 of these isolates were confirmed to be resistant to LFX by using 2.0 mg/l as cut-off value, and the remaining 15 isolates were susceptible to LFX. The LFX MICs for MTB isolates with the mutation Ala90Val ranged from 2.0 to 8.0 mg/l. In addition, the lower LFX values were observed for Asp94Ala mutants, ranging from 1.0 to 4.0 mg/l; however, all five other *gyrA* mutants had MICs of ≥4.0 mg/L. At the same time, the 2 genetically heterogeneous populations harbored MIC values of 4.0 mg/L. Importantly, all 13 LFX-susceptible isolates with *gyrA* mutations belonged to Ala90Val and Asp94Ala group.

## DISCUSSION

Fluroquinolones are widely used in the treatment of MDR-TB, and have also been contained in several regimen trials for the treatment of drug-susceptible and drug-resistant TB (10, 18), highlighting the urgent need for early diagnosis of *in vitro* FQ susceptibility. In the present study, our data demonstrated that more than one quarter of LFX-resistant TB cases identified by molecular diagnostics were phenotypically LFX-susceptible. The high rate found in this study may be majorly due to the fact we used broth-based assays for drug susceptibility testing instead of the solid agar-based methods, which are more likely to identify the drug-resistant MTB isolates with low-level resistance (19). In addition, although phenotypical DST is considered as gold standard for detection of *in vitro* drug susceptibility, there is increasing evidence that the low-level resistance is associated with a marked decrease in reproducibility of this phenotypical method, even in the sophisticated laboratories (8, 20). As a consequence, the isolates with MICs that is very close or equal to the defined critical concentration for the tested drug are prone to be missed. These isolates are termed “challenging isolates”, as they are responsible for the majority of discordant results between phenotypical and genotypical method.

In addition, the heteroresistance is another important cause of the discordant LFX susceptibility results (21). On one hand, the coexistence of wide-type and mutant tubercle bacilli may result in a shift toward lower MIC value compared with that of mutant bacilli (16). Despite having the mutant population (Asp94Gly) with high-level resistance to LFX, the MIC values of two isolates located around the critical concentration. On the other hand, the Sanger sequencing failed to identify the presence of mutant tubercle bacilli from the two isolates recovered from clinical sputum. Numerous previous studies have demonstrated that the *gyrA* mutations do not only seem to have an impact on the FQ resistance but also on the fitness of the strains (22). Thus, the loss of relative fitness associated with specific mutations in *gyrA* gene may explain why the bacterial population structure changed significantly during the subculture period.

It is well known that various *gyrA* mutations are associated with different levels of resistance to LFX (11). In consistent to previous findings, we found that the MICs of isolates with either Ala90Val or Asp94Ala mutation are scattered around the critical concentrations of LFX (11, 23). In a previous retrospective study, the treatment failure with FQ-containing regimen was reported in patients infected with isolates harboring these mutations, suggesting that they are probable resistant to FQ (24), but may be missed by the conventional DST methods. Considering that Ala90Val and Asp94Ala mutation are the most predominant mutation sites conferring FQ resistance in China (11), the high prevalence of discordant levofloxacin resistance could be expected in clinical practice.

Our findings provide important implications for the diagnosis and management of drug-resistant tuberculosis. First, the low-level drug resistance is believed to be the most important reason for the disputed LFX resistance. In other words, the molecular diagnostic algorithm outperforms the phenotypical DST methods for detecting the LFX-resistant isolates with mutations conferring proximity of MICs to the critical concentration. Thus, the reliable LFX-resistance by molecular methods should be adopted in individualization of multidrug regimens. Second, operational aspects of the methods should be mentioned. Conventional DST methods remain technically cumbersome, and the culture-based characteristic requires time to obtain positive isolates (3). In contrast, the commercial molecular DST methods could be directly used on sputum samples (preferably smear positive), which saves intervals over phenotypic methods requiring culture. A recent external quality assessment (EQA) study of TB laboratories in China demonstrated that approximate 40% of laboratories failed to provide certified conventional DST results; however, the EQA results for molecular diagnostics were satisfactory for these laboratories (25). Taken together, these results highlight the potential clinical application of molecular diagnostics in diagnosis of drug-resistant TB, especially for resource-limited settings. Third, previous molecular epidemiological studies on FQ resistance always included phenotypically resistant MTB isolates determined by phenotypical DST (11, 23). In view of their restricted capability to detect low-level resistant isolates, the prevalence of mutations within *gyrA* locus conferring low-level resistance may be underestimated.

We also acknowledge the limitations in the study. First, the major limitation is the small sample size in our analysis. The first commercial kit for detecting FQ susceptibility, MeltPro TB assay, was approved by Chinese FDA in 2018. Hence, the short-term application of this method become the major barrier to recruit more cases. Further study will be carried to validate our conclusion in larger samples. Second, the discordant LFX susceptibility was not assessed by another phenotypic DST method. It is difficult to compare the capability of different phenotypic methods to detect low-level LFX resistance. Third, the clinical outcomes of these cases with discordant LFX susceptibility were not evaluated due to the high proportion of loss to follow-up.

In conclusion, our data demonstrate that more than one quarter of LFX-resistant isolates by molecular method are susceptible to LFX using the phenotypic DST. The high prevalence of discordant LFX resistance is majorly due to the isolates with specific mutations within *gyrA* gene conferring proximity of MICs to the critical concentration. Our results will provide important implications into the clinical interpretations of discordant LFX resistance between phenotypical and genotypical DST methods.

## ACKNOWLEDGEMENTS

This work was supported by the Tongzhou District Science and Technology Committee (KJ2019CX016), the Beijing Talents Foundation (2017000021223ZK39), the Beijing Municipal Administration of Hospitals’ Youth Programme (QML20171601), and Beijing Municipal Administration of Hospitals Incubating Program (PX2019061).

